# scBERT as a Large-scale Pretrained Deep Language Model for Cell Type Annotation of Single-cell RNA-seq Data

**DOI:** 10.1101/2021.12.05.471261

**Authors:** Fan Yang, Wenchuan Wang, Fang Wang, Yuan Fang, Duyu Tang, Junzhou Huang, Hui Lu, Jianhua Yao

**Author notes:** These authors contributed equally: Fan Yang, Wenchuan Wang, Fang Wang. These authors jointly supervised this work: Hui Lu, Jianhua Yao. Hui Lu, Jianhua Yao.

## Abstract

Annotating cell types based on the single-cell RNA-seq data is a prerequisite for researches on disease progress and tumor microenvironment. Here we show existing annotation methods typically suffer from lack of curated marker gene lists, improper handling of batch effect, and difficulty in leveraging the latent gene-gene interaction information, impairing their generalization and robustness. We developed a pre-trained deep neural network-based model scBERT (single-cell Bidirectional Encoder Representations from Transformers) to overcome the challenges. Following BERT’s approach of pre-train and fine-tune, scBERT obtains a general understanding of gene-gene interaction by being pre-trained on huge amounts of unlabeled scRNA-seq data and is transferred to the cell type annotation task of unseen and user-specific scRNA-seq data for supervised fine-tuning. Extensive and rigorous benchmark studies validated the superior performance of scBERT on cell type annotation, novel cell type discovery, robustness to batch effect, and model interpretability.

---

Single-cell RNA sequencing (scRNA-seq) has been extensively used for the characterization of complex tissues and organisms at single-cell level^1–3^, which has revolutionized transcriptomic studies. Accurate cell type annotation on scRNA-seq is critical for biological and medical researches^4^. Cell type annotation methods can be categorized into three types: annotation using marker genes, correlation-based methods, and annotation by supervised classification^5^.

Cluster-then-annotate is the commonly-used manner^6^ where manually curated marker genes identified from literatures are employed to assign cell types for clusters derived from unsupervised learning^5^. However, selecting the marker genes depends on researchers’ prior knowledge, and is therefore prone to bias and errors^7^. Additionally, maker genes for interested cell types are not always available, and novel cell types don’t have marker gene sets yet. Besides, most cell types are determined by a set of genes instead of a single marker gene^8^. Without a proper method to integrate the expression information of multiple marker genes, it is difficult to guarantee a unified and accurate cell type assignment for each cluster^9,10^. For example, some automatic annotation methods are built upon the hypothesis that marker genes should have high expression in cells. However, even some well-documented marker genes do not have high expression in all the cells in the corresponding cell types^11^. Therefore, the absence or fluctuation of the expression of these marker genes might significantly affect the preciseness of marker-gene-based methods.

Instead of relying on a spot of marker genes, correlation-based methods measure the correlation of gene expression profiles between the query samples and reference dataset^5^. These methods are potentially affected by the batch effect across platforms and experiments^12^. Although there exist batch-effect correction methods, it’s still challenging to distinguish true biological diversity from technical differences and thus preserve important biological variations^13^. Meanwhile, the commonly-used similarity measures (i.e. cosine similarity, Spearman’s correlation, and Pearson correlation) may not be robust or efficient for measuring the distance between two sets of high-dimensional sparse scRNA-seq data^14^.

Annotation by supervised/semi-supervised classification mehods follows the classic paradigm in machine learning (ML) that recognizes patterns in the gene expression profiles and then transfers the labels from labeled datasets to unlabeled datasets^5^. They are widely used recently due to their robustness to noise and variability of data, and their independence of artificially selected marker genes. Nevertheless, most of these methods need to perform highly-variable-gene (HVG) selection and dimensionality reduction before inputting the data into the classifier due to their limited model capacity^15–19^. However, HVGs are variable across different batches and different datasets, hindering their generalization ability across cohorts^16^. Dimensionality reduction like PCA may lose high-dimensional information as well as gene-level independent interpretability. Furthermore, the parameter settings of HVG selection and PCA in these methods are far to reach a consensus and inevitably introduce artificial bias for performance evaluation^15–19^. Given that the HVGs are selected based on the expression variance across the whole dataset where the dominant cell types account for the most variance, there is a risk of overlooking the key genes of rare cell types. Selecting HVGs ignores co-occurrence and biological interaction of genes (especially HVG and non-HVG), which are useful for cell type annotation^20^. Besides, simple classifiers such as fully connected networks couldn’t efficiently capture gene-gene interactions. Therefore, a new method with improved pattern recognition ability is required to overcome the above issues of under-fitting to large-scale datasets.

Recently, a growing number of deep learning-based methods have been applied to scRNA-seq data analyses and achieved superior performance^21–23^. BERT (the Bidirectional Encoder Representations from Transformers) is a state-of-the-art (SOTA) Transformer-based language representation learning model. It has made breakthrough progress in the fields of natural language processing (NLP) due to the powerful self-attention mechanism and long-range information integration capability introduced by Transformer layers^24,25^. BERT’s paradigm of pre-training and fine-tuning enables the use of large-scale unlabeled data to improve the generalizability of AI model. Inspired by such exciting progress, we developed scBERT (*s*ingle-*c*ell *B*idirectional *E*ncoder *R*epresentations from *T*ransformers) model for the cell annotation task of scRNA-seq data. Following the pre-training and fine-tuning paradigm, we validated the power of applying self-supervised learning on large-scale unlabeled scRNA-seq data to improve the model’s generalizability and overcome the batch effect. Extensive benchmarking indicated that scBERT can provide robust and accurate cell type annotations with gene level interpretability. To the best of our knowledge, scBERT pioneered the application of Transformer architectures in scRNA-seq data analysis with innovatively designed embeddings for genes.

## Results

### The scBERT algorithm

The original BERT^25^ proposed a revolutionary recipe that generates a generic knowledge of language by pre-training and then transferring the knowledge to downstream tasks of different configurations using fine-tuning. Following BERT’s mentality and paradigm, we developed a novel and unified architecture named scBERT (Fig.1), which learns a general scRNA-seq knowledge by being pre-trained on millions of unlabelled scRNA-seq data with a variety of cell types from different sources, and assigns cell types by simply plugging in a classifier and fine-tuning the parameters supervised by reference datasets. Pre-training enables the model to learn general syntax of gene-gene interactions, which helps to remove the batch effects across datasets and improve the generalizability (Extended Data Figure 1a). Fine-tuning ensures that the output embedding for each gene encodes context information that is more relevant to the transcriptional profiles of the reference dataset. To annotate a query cell, scBERT computes the probability for the cell to be any cell type labeled in the reference dataset by mining the high-level implicit patterns (Extended Data Figure 1b). Of note, if there is no cell type to assign with high confidence, the query cell would be labeled as ‘unassigned’ to prevent incorrect assignment and to allow novel cell type discovery. The scBERT model has some innovative designs compared to the original BERT model to unleash the power of BERT in the cell type annotation task.

**Figure 1.**
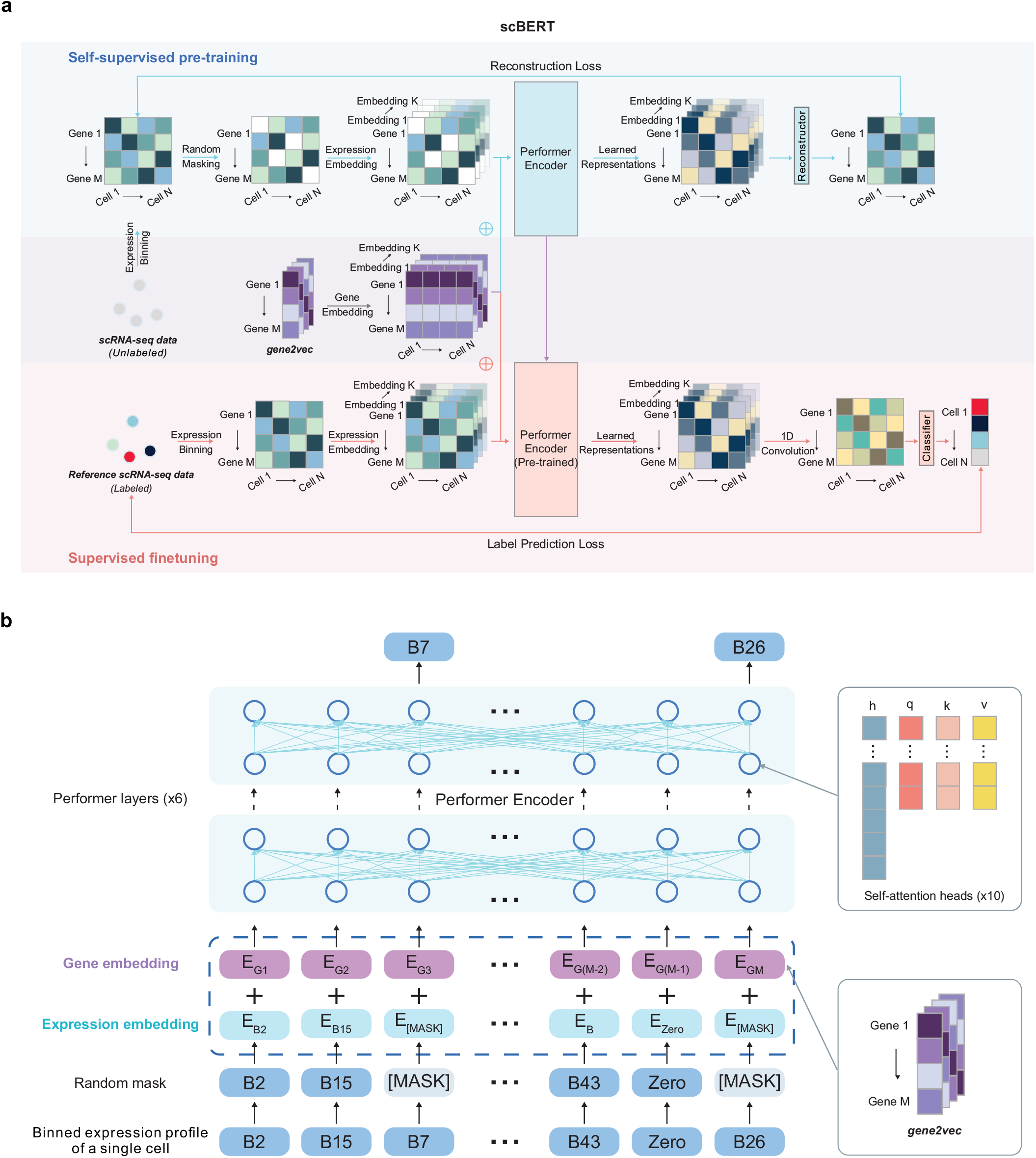
Overview of the scBERT model. **a,** Self-supervised learning on unlabeled data and fine-tuning on task-specific data. At the self-supervised pre-training stage, unlabeled data were collected from PanglaoDB. Masked expression embedding and gene embedding were added as input and then fed into the Performer blocks. The reconstructor was used to generate outputs. Outputs for masked genes were used to calculate the reconstruction loss. At the supervised fine-tuning stage, the task-specific scRNA-seq data were input into the pretrained encoder. The output representation then passed a 1-dimensional convolution layer and a classifier to generate the cell type prediction. ⊕: element-wise addition. The Performer encoder is the component that is shared between the models used in the pre-training and fine-tuning stages. The reconstructor and the classifier are independently and separately employed for the models during the pre-training and fine-tuning processes. **b,** Illustration of the embeddings of scBERT. The preprocessed scRNA-seq data are first converted into discretized expression, and then the non-zero expressions are randomly masked. Taking the first gene as an example, the gene embedding E_G1_ (represents the gene identity from gene2vec falling into the first bin) and the expression embedding E_B2_ (represents the gene expression falling into the second bin and being transformed to the same dimension as the E_G1_) are summed and fed into scBERT to generate representations for genes. The representations are then used for pre-training or finetuning.

First, the embedding of BERT includes token embedding and position embedding^25^. Our design of embeddings is similar to BERT in some aspects while having unique features to leverage gene knowledge. The token embedding of the original BERT is a discrete variable (standing for a word), while the raw expression input to our model is a continuous variable (standing for the expression of a gene in a single cell) with biological and technical noise. We draw on the bag-of-words technology in the NLP^26^ field to bin the expressions of genes (which could be considered as the gene transcript frequency in each cell), thus converting them to discrete values with additional benefits of the reduction of the data noise to some extent. Since shuffling the columns of our input doesn’t change its meaning (like the extension of BERT to understand tabular data with TaBERT^27^), absolute positions are meaningless for gene. Instead, gene embeddings were obtained from the gene2vec^28^ to represent the gene identity (each gene has a unique gene2vec embedding) and could be viewed as a relative embedding^26^ to capture semantic similarity from the aspect of general co-expression. The co-expression genes retain closer representations, and distributed representation of genes has been proved to be useful for capturing gene-gene interactions^28^. In this way, scBERT formalizes information on the gene expressions for the Transformer efficiently and generates single-cell specific embedding (abbr. scBERT embedding) that represents the cell-specific expression (Extended Data Figure 1c) after pre-training.

Second, existing single-cell methods have to preprocess the raw data with selection or manipulation of genes (i.e., HVG selection, manually selecting marker genes, and PCA) due to the limited model capability on efficiently modeling high-dimension data^9,10,29–32^. They would unavoidably bring artificial bias and overfitting problem, which in turn may severely impair their generalizability. Conversely, Transformer with large receptive filed could effectively leverage the global information possessed in scRNA-seq data and learn comprehensive global representation for each cell by unbiasedly capturing long-range gene-gene interaction. Due to the computational complexity, the input sequence of Transformer is limited to the length of 512, while most of the scRNA-seq data contain over 10,000 genes. Therefore, we replaced the Transformer encoder used in BERT with Performer^33^ to improve the scalability of the model to tolerate over 16,000 gene inputs. With Performer, scBERT keeps the full gene-level interpretation, abandons the use of HVGs and dimensionality reduction, and lets discriminative genes and useful interaction come to the surface by themselves (Extended Data Figure 1d). Thus, scBERT allows for the discovery of gene expression patterns and longe-range dependency for cell type annotation in an unbiased data-driven manner. scBERT is stable and robust, instead of relying heavily on the hyperparameter selection (Extended Data Figure 1e).

### Evaluating cell type annotation robustness on intra-dataset

We first benchmarked the performance of scBERT against comparison methods on 9 scRNA-seq datasets, covering 17 major organs/tissues, more than 50 cell types, over 500,000 cells, and mainstream single-cell omics technologies (i.e., Drop-seq, 10X, SMART-seq, and Sanger-Nuclei), comprehensively considering the diversity of data size and data complexity^34^ (Supplementary Table 1). For comparison, marker-gene-based methods (SCINA, Garnett, scSorter), correlation-based methods (Seurat v4, SingleR, scmap_cell, scmap_cluster, Cell_ID(c), Cell_ID(g)), and ML-based methods (SciBet, scNym) were used (Supplementary Table 2). For each of the datasets, we applied the 5-fold cross-validation strategy to avoid the influence of random results on the conclusion. scBERT surpassed the performance of comparison methods in both accuracy and macro F1 score on most of the datasets (Fig. 2a, Extended Data Figure 2). Among the intra-dataset, Zheng68K dataset from human peripheral blood mononuclear cells (PBMC) is the most representative dataset for benchmarking cell type annotation methods. Due to the severe cell type imbalance and the extremely high similarity between subtypes, even the SOTA method could not achieve an accuracy above 0.71. The performance of scBERT with completely deletion of reported marker genes is already on par with the best performance of existing methods (Extended Data Figure 1b), demonstrating the superior of scBERT’s pattern recognition ability on gene expressions compared with those methods heavily depending on known marker genes. With the addition of marker genes, scBERT could capture more comprehensive gene expression patterns constructed by marker genes. With all genes as inputs, scBERT significantly surpassing SOTA methods with a large margin (Fig. 2b-c, Extended Data Figure 3, Extended Data Figure4a) (scBERT F1 score=0.691, accuracy=0.759; best F1 score of other methods=0.659, accuracy=0.704) on overall cells, and achieved the highest performance (F1 score=0.788 vs. 0.617, P-value=9.025e^−5^;accuracy=0.801 vs. 0.724, P-value=2.265e^−5^) for CD8+ cytotoxic T cells and CD8+/CD45RA+ T cells that are highly similar and hard to distinguish in previous studies^35^. The results indicated that scBERT could recognize underlying gene expression patterns and long-range gene-gene dependency after pre-training, capture diverse feature subspace by multi-head attention and enjoy comprehensive high-level representation of cell-type-specific global information.

**Figure 2.**
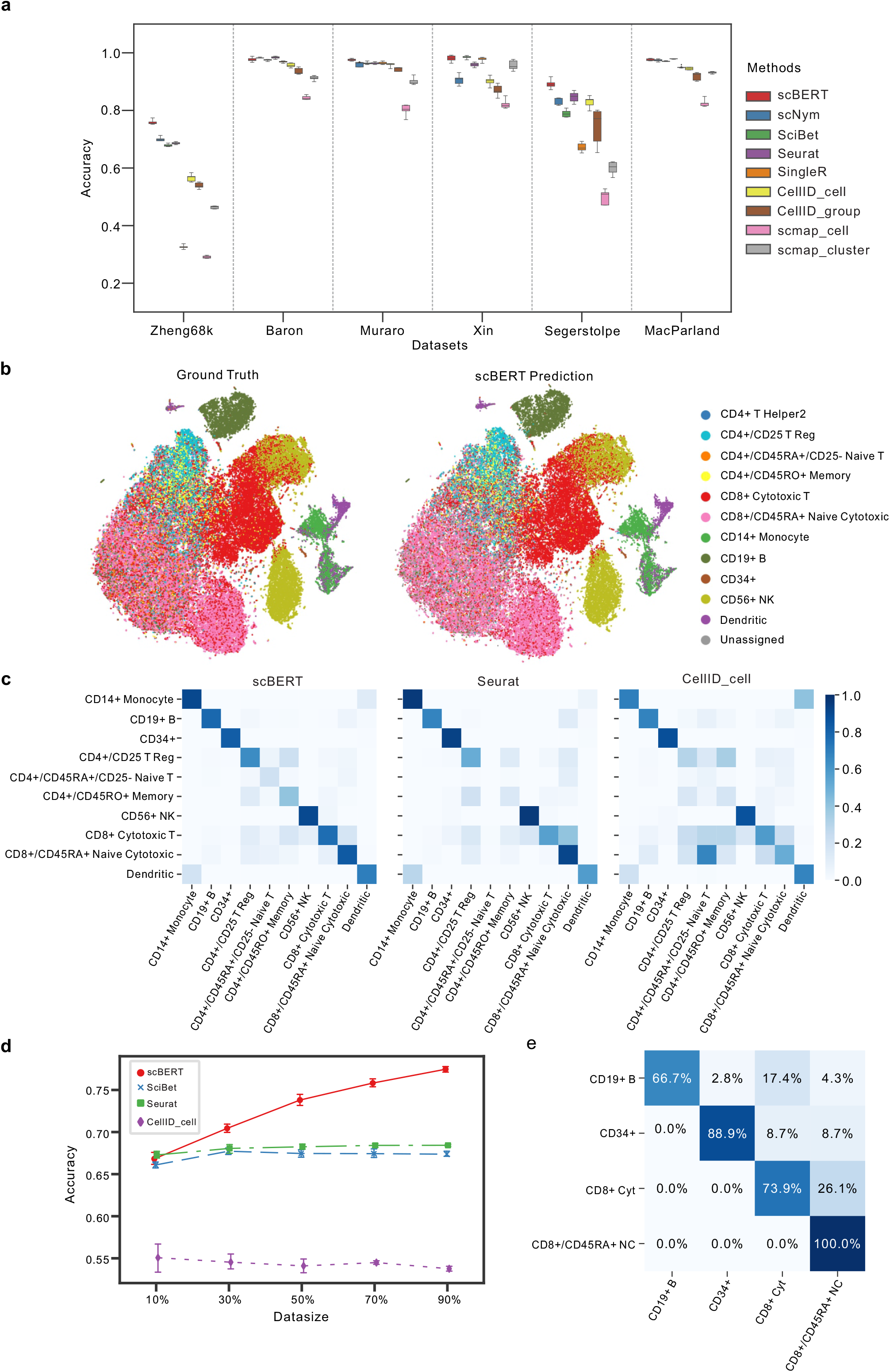
Benchmarking and robustness evaluation by intra-dataset cross-validation. **a,** Performance of cell type annotation methods measured by accuracy and F1 score on n=9 datasets using 5-fold cross-validation. Box plots show the median (center lines), interquartile range (hinges), and 1.5 times the interquartile range (whiskers). The accuracy of the Zheng68K, Baron, Muraro, Xin, Segerstolpe and MacParland datasets is shown from left to right. The F1 score of these datasets is shown in Extended Data Figure 2a. The performance of SCINA, Garnett and scSorter is shown in Extended Data Figure 2b. The results of Tucker dataset, Lung dataset and Human Cell Atlas dataset are shown in Extended Data Figure 2c-d. **b,** t-SNE plot of the whole Zheng68K dataset (n= 68,450 cells). Left panel is colored by expert-annotated cell types from the original research, right panel is colored by scBERT prediction results. The t-SNE plots of the annotation of comparison methods are shown in Extended Data Figure 3. **c,** Heatmaps for the confusion matrices of the cross-validation results on the Zheng68K dataset for scBERT, Seurat and CellID_cell. The confusion matrices of other methods are included in Extended Data Figure 4a. **d,** The influence on the cell type annotation performance by splitting different proportions of the Zheng68K dataset as the reference set for fine-tuning. The standard deviations are shown as the error bar. **e,** Heatmap for the confusion matrices of scBERT of cross-validation on the imbalanced dataset reconstructed from Zheng68K dataset. The confusion matrices of other methods are included in Extended Data Figure 4b. The detailed reconstruction process is introduced in the method section.

Notably, the list of the top-tier methods changes across different tasks and datasets while scBERT is always among it. For instance, the top-tier methods on inter-dataset (i.e., scNym and Seurat) performed badly on Xin dataset in Fig.2. These uncertainties in performance reflect the limitations of the comparison methods in their generalizability, as well as the generalization of our method across all the benchmarking datasets.

To explore whether the number of cells of a reference dataset affects the performance of scBERT, we constructed a series of reference datasets from the Zheng 68K dataset by uniformly subsampling it proportionally from 10% to 90% (Fig. 2d). With only 30% cells from Zheng 68K dataset), scBERT beated all comparison moethods, and its performance improved rapidly as reference cell number increased.

We next tested the robustness of scBERT when the distributions of cell types were severely biased. Four cell types from the Zheng 68K dataset (CD8+ cytotoxic T cells, CD19+ B cells, CD34+ cells, CD8+/CD45RA+ naive cytotoxic cells) with transcriptomic similarity between each pair, were selected for class-imbalanced tests. scBERT surpassed all comparison methods (accuracy=0.840 and F1 score=0.826). Seurat misidentifed CD8+ cytotoxic T cells to CD8+/CD45RA+ naive cytotoxic cells and SingleR misclassified all CD19+ B cells due to their rarity, while scBERT performed the least misclassification rate even though high similarity exists between the two cell populations (Fig. 2e, Extended Data Figure 4b). Overall, the results indicate that scBERT is robust to class-imbalanced datasets.

### Cell type annotation across cohorts and organs

In real-world circumstances, the reference dataset and query dataset are always sourced from multiple studies, and even different sequencing platforms, where the batch effects can lead to poor performance on cell type annotation (Fig.3a). Here, we benchmarked scBERT and comparison methods by employing a leave-one-dataset-out strategy with human pancreas datasets generated by distinct sequencing techniques (Baron^36^, Muraro^37^, Segerstolpe^38^, and Xin^39^) (Fig.3, Extended Data Figure 5). ML-based methods (scBERT, scNym, and SciBet) achieved the best results, indicating that cell-type-specific patterns could be discovered by pattern recognition without being affected by batch effects, while Seurat relies on compulsive batch correction before the annotation. For cross-cohort data, scBERT achieved a superior performance with accuracy of 0.992 compared to scNym (accuracy=0.904) with a large margin (about 10%)and outperformed other popular methods (accuracy: SciBet 0.985, Seurat 0.984, SingleR 0.987) (Fig.3b). scBERT correctly annotated most cells (> 97%) in the Muraro dataset and over 99% in the other three datasets, demonstrating the superb and stable performance of our method in cross-cohort tasks. In the contrast, scNym misclassified the alpha cells as beta cell type and was confused with the beta and delta cells (Fig.3e-f). We then used cells from different organs to benchmark the performance of scBERT and comparison methods on cross-organ dataset. The experiment results demonstrated that scBERT is on par with comparison methods on cross-organ task (Extended Data Figure 5b). scBERT showed its robustness in identifying cells from different sequencing technologies, experiments, different disease states (type 2 diabetes (T2D) and health), and even different organs.

**Figure 3.**
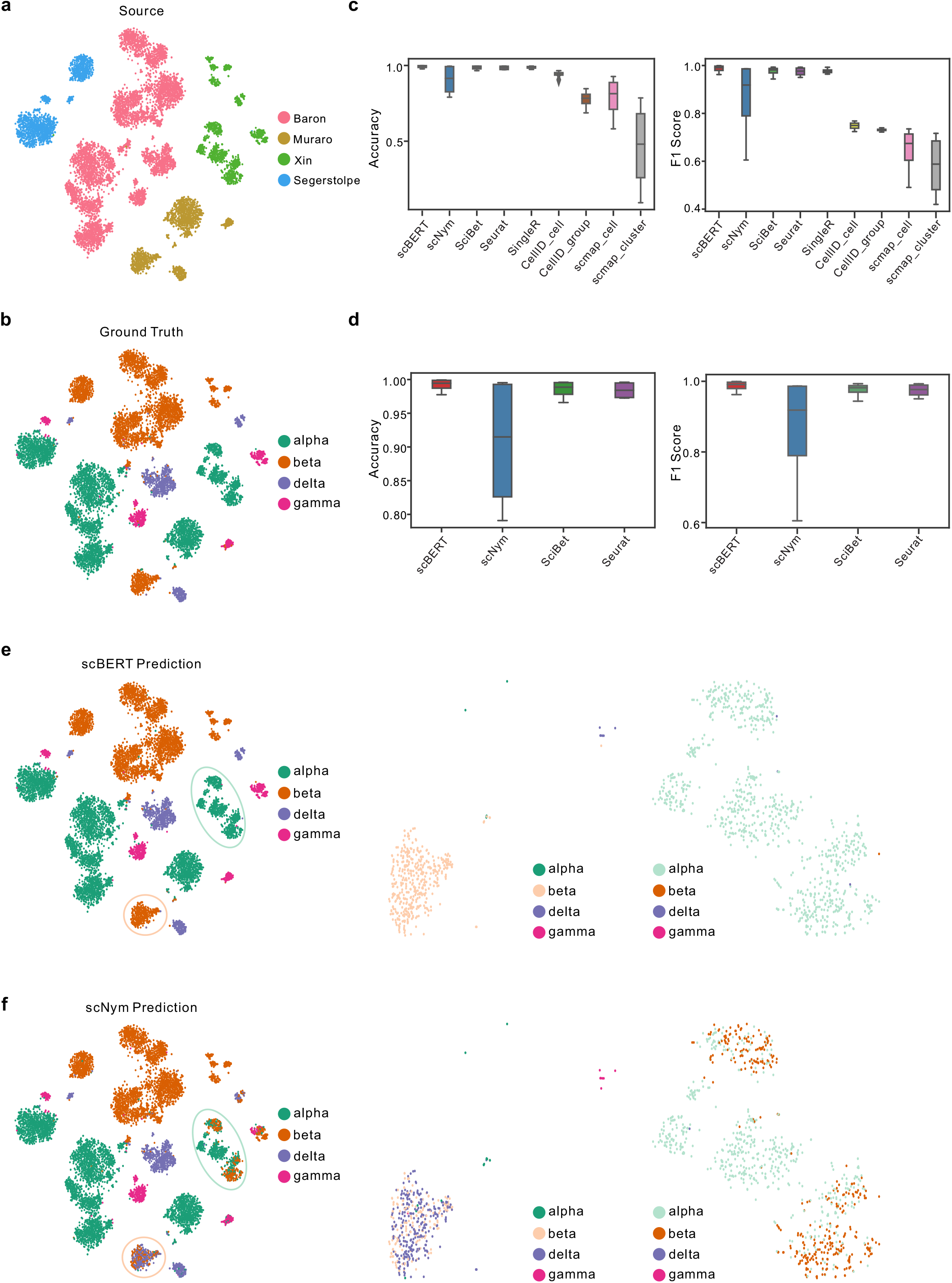
Performance of scBERT across independent datasets generated by different single-cell sequencing technologies. **a,** t-SNE representation of 10,220 cells from four independent datasets (Baron, Muraro, Segerstolpe, and Xin) generated by different sequencing platforms (inDrop, CEL-Seq2, SMART-Seq2, and SMARTer). Cells are colored by the source of datasets. **b,** t-SNE representation of alpha, beta, delta, and gamma cells from four pancreas datasets colored by the annotated cell types provided by the atlas from the original paper. **c,** Comparison of accuracy and F1 score of inter-dataset cross-validation among different methods. The lower and upper hinges denote the first and third quartiles, with the whiskers in the range of 1.5 times the interquartile. **d,** Zoomed-in plot of accuracy and F1 score of the top tier methods. The lower and upper hinges denote the first and third quartiles, with the whiskers in the range of 1.5 times the interquartile. **e,** t-SNE representation of alpha, beta, delta, and gamma cells from four pancreas datasets (left), beta cells from Muraro dataset (middle), and alpha cells from Segerstolpe dataset (right) colored by scBERT prediction. **f,** t-SNE representation of alpha, beta, delta, and gamma cells from four pancreas datasets (left), beta cells from Muraro dataset (middle), and alpha cells from Segerstolpe dataset (right) colored by scNym prediction. t-SNE plots of other comparison methods are shown in Extended Data Figure 5a.

### Discovery of novel cell types

The reference dataset may not cover all cell types present in the query dataset in most tasks. The marker-based methods are hindered by the manually selected marker of known cell types and might be difficult to distinguish unseen cell types, while the correlation-based methods usually force the model to assign a novel class to a closest known class. The ML-based methods could automatically and actively detect the novel cell types by checking the predicted probability. Besides, scBERT enjoys potential advantages. First, the multi-head attention mechanism allows scBERT to extract information from different representation subspaces, which might benefit for capturing the subtle difference between novel cell type and known cell types. Second, during pre-training on large-scale diverse data set, scBERT possibly has seen the novel cells and learnt their unique patterns. Third, Transformer with large receptive filed could effectively learn comprehensive global representation by capturing long-range gene-gene interaction, which may better characterize and distinguish novel cells. ^41^ Not surprisingly, scBERT performed the best on novel cell types and achieved the first-class performance on the known cell types (Fig.4). CellID_cell performed well on known cell types, while failed to discover any novel cells. SciBet and scmap_cluster is prone to assigning “unknown” labels to those cells from known types, which greatly reduce their accuracy of known cell type classification. Compared to SciBet and scmap_cluster, our method achieves the superior accuracy on both the novel class (scBERT=0.329 vs. SciBet=0.174 and scmap_cluster=0.174) and known classes (scBERT=0.942 vs. SciBet=0.784 and scmap_cluster=0.666). Taken together, these results suggested that scBERT can correctly discover novel cell types that are not present in original reference datasets and remain accurate prediction performance of other cell types.

**Figure 4.**
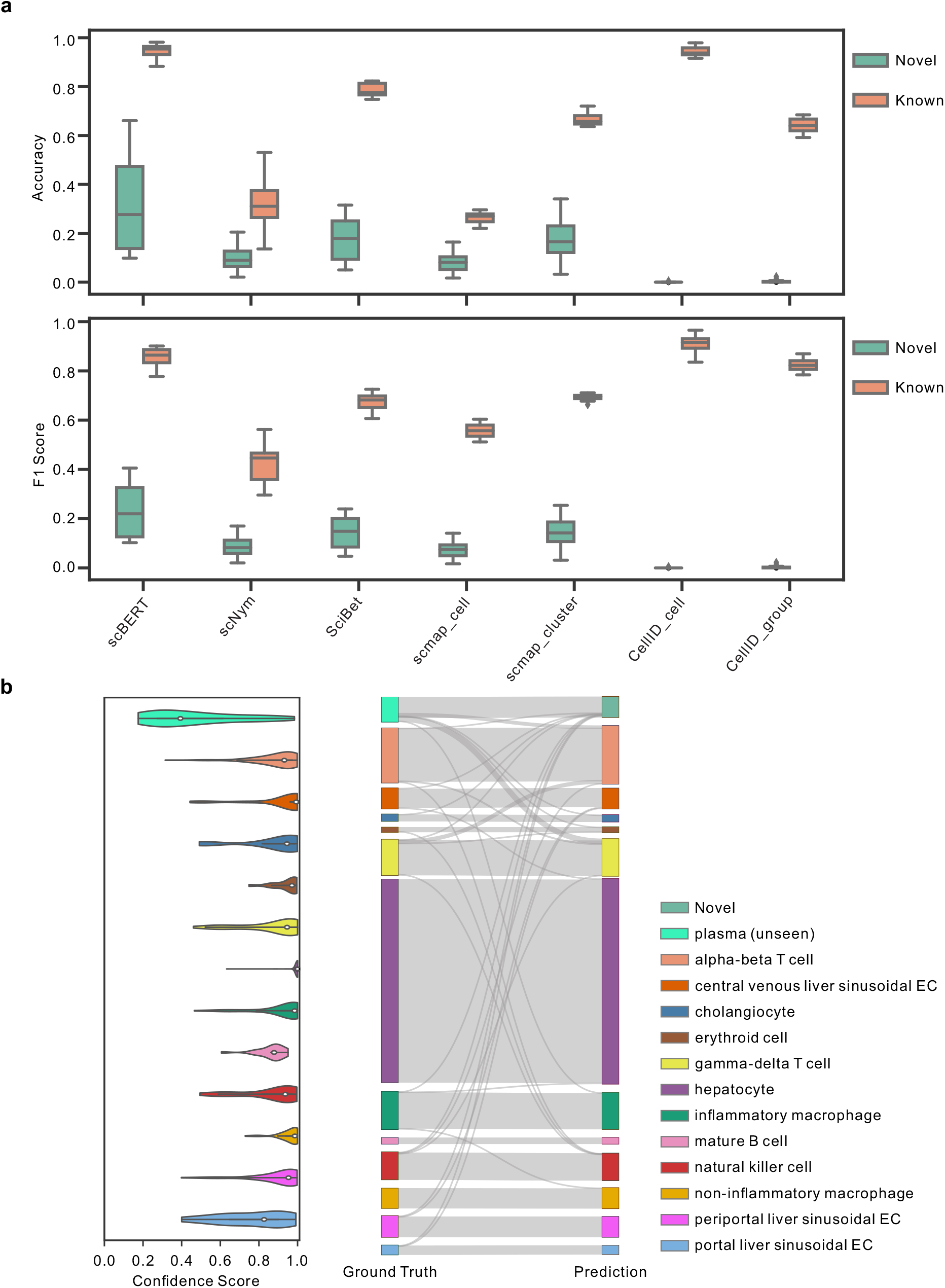
Identification of novel cell types. **a,** Performance of scBERT on the MacParland dataset from human liver tissue by removing alpha-beta T cell, gamma-delta T cell, mature B cell, and plasma cell populations respectively during the scBERT training process. The accuracy and F1 score of both novel cell types and known cell types are shown in the boxplots. Box plot shows the median (center lines), interquartile range (hinges), and 1.5 times the interquartile range (whiskers). **b,** The confidence scores provided by scBERT for the cell types of MacParland (left), the cells with low probability of model prediction (probability<0.5) for all known cell types are assigned as potential novel cell types. Sankey plot comparing scBERT predictions on known and novel cell types with original cell-type annotations for the MacParland dataset (right), where plasma cells are labeled as novel cell type since they are unseen by the scBERT training process.

### Investigating scBERT model interpretability

Existing ML methods select HVG or reduce dimension due to their simplified network architecture and low model capacity hence destroying the gene-level interpretability. In contrast, the attention mechanism employed in scBERT naturally provides hints for the decision-making of the model using every individual gene.

Here, we took the Muraro dataset as an illustration and top-attention gene lists were produced for the four kinds of pancreas islet cells, with well-studied biological functions (Fig.5a). The top-attention genes included reported markers of specific cell types (i.e., LOXL4 for alpha cell and ADCYAP1 for beta cell^40^)(Extended Data Figure 6a). Almost all the top-attention genes except markers were identified as differentially expressed genes using DESeq^41^, as potential novel markers (Fig. 5c, Extended Data Figure. 6b). For instance, SCD5 hasn’t been reported as a cell-type-specific marker for beta cells, but in a GWAS study, a novel loci for T2D susceptibility was fine-mapped to a coding variant of SCD^42^. The results demonstrated that scBERT could facilitate understanding the cell type annotated and provide some support for further biological findings.

**Figure 5.**
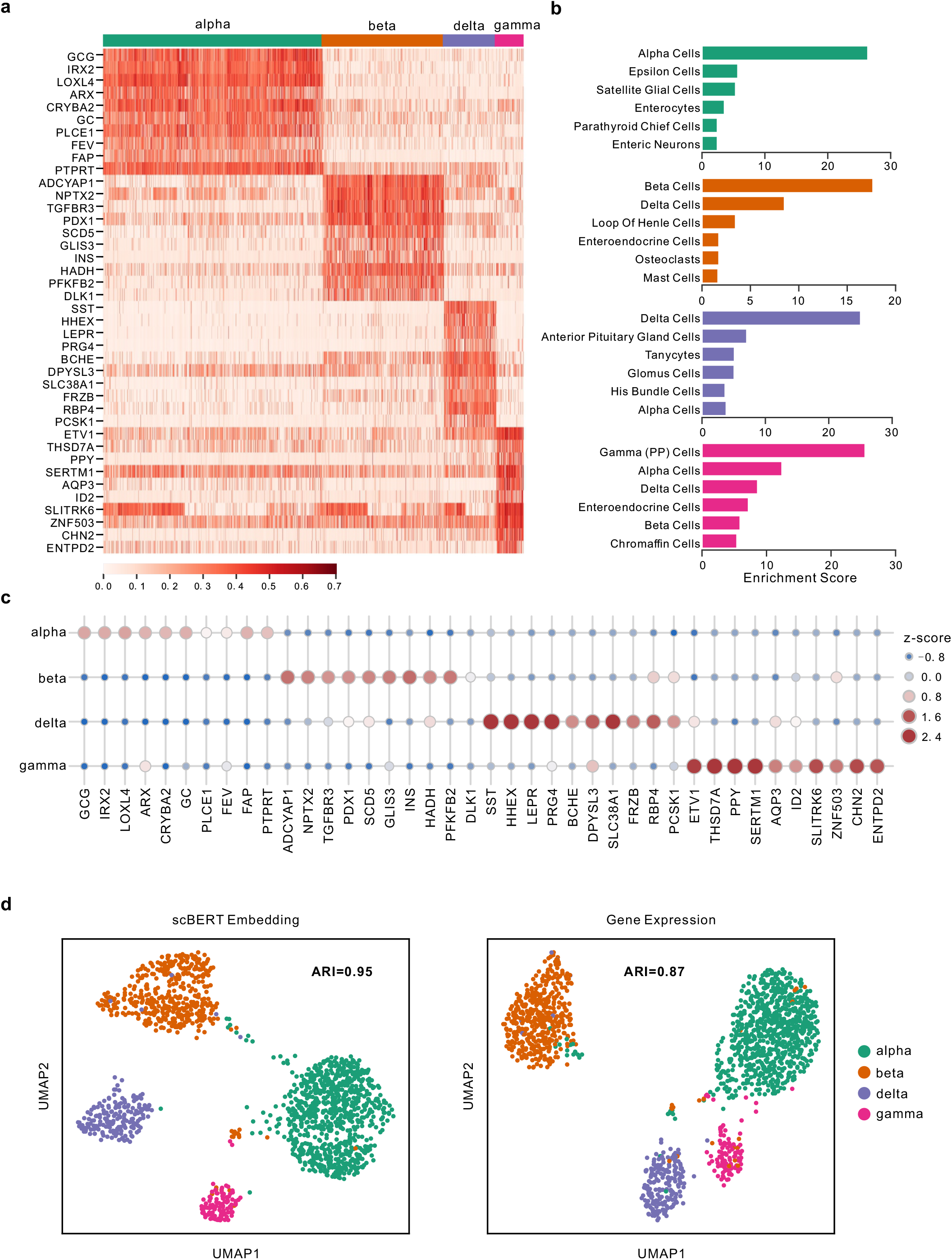
Model interpretability. **a,** Heatmap for the attention weights provided by scBERT on Pancreas cell type annotation task. The detailed attention estimation process is described in Method section. Top 10 genes with highest attention weights are listed for each cell type. Complete top gene list is in Supplementary Table 3. **b,** The results of enrichment analysis of the top attention genes from scBERT, with the complete information in Supplementary Table 4-15. **c,** Dotplot showing z-scores among the 10 genes receiving highest attention and the cell types. As shown in the legend, the size and color of each dot reflect the z-score. **d,** UMAP representation of alpha, beta, delta, and gamma cells from Muraro dataset colored by cell types. Left UMAP is based on the scBERT embedding of each cell, while right UMAP is based on the raw expression of each cell. The ARI score is calculated and shown in the plot.

Enrichment analysis was performed for the top 50 attention gene lists using various gene-set libraries and the results revealed there were some interesting relationships between top enriched terms and the corresponding cell types (Fig.5b, Supplementary Table 3-15). In particular, with the cell-type-associated gene-set library from PanglaoDB, the top-one-enriched term for each type always hits the true cell population. As another example, insulin secretion and AMPK signal pathway, the top-two-enriched KEGG pathways in beta cells, are vital to beta cell function. Furthermore, the scBERT embedding is more distinguishable for cell type annotation than the raw gene expression based on the clustering performance (ARI: 0.95 vs. 0.87), indicating the efficiency of scBERT in learning single-cell-specific representation, which can be used for downstream analysis (Fig.5d).

## Discussion

In order to improve the generalization ability of the cell type annotation algorithm and the interpretability of individual gene importance, we developed scBERT, a deep learning model with a multi-head attention mechanism and self-supervised strategy, to learn domain-irrelevant gene expression patterns and interaction from the whole genome expression of large-scale unlabeled scRNA-seq data, transfer the general knowledge to cell type annotation task by fine-tuning, and trace back to the importance of each individual gene for model interpretability. By systematically analyzing the components of scBERT, we gain several insights of the application of Transformer in single-cell data analysis (i.e., benefit of pre-training, recognization of non-marker pattern, detection of subtle gene-gene interaction, single-cell specific embedding and hyperparameters sensitivity) (systematic analysis in Method, extended data figure 1).

scBERT surpasses the existing advanced methods on diverse benchmarks collectively involving 9 single-cell datasets, 17 major organs/tissues, more than 50 cell types, over 500,000 cells, and the mainstream single-cell omics technologies (i.e., Drop-seq, 10X, SMART-seq, and Sanger-Nuclei), indicating its generalization and robustness. Notably, we employed the accuracy, macro-F1 score, and confusion matrix as evaluation metrics for benchmarking the performance of cell type annotation methods on their classification ability for a fair comparison in this study.

To the best of our knowledge, there is currently no research on applying Transformer architectures to gene expression data analysis. The originally designed end-to-end scBERT framework with gene expression embedding and self-learning strategy has superior performance, interpretability, and generalization potential on cell type annotation task. Beyond that, scBERT can also be migrated to other tasks by simply modifying output and supervision signals. scBERT, as an effective cell type annotation tool, has been released on the platform for public usage. We hope that scBERT could improve understanding of cell-type-associated gene-gene interaction and nurture the revolution of AI paradigm in single-cell RNA-seq analysis.

Despite the above advantages, the scBERT may face potential limitations including gene expression embedding, modeling gene interactions, and the masking strategy during the pre-training stage.

First, the token embedding of the original BERT is for discrete variable (standing for a word), while the expression input is a continuous variable (standing for the expression of a gene in a single cell), which may have biological and technical noise. scBERT converts them to discrete values and could thus reduce some data noise compared with existing methods which utilize the expression values directly. However, it sacrifices some data resolution and there is still room for optimizing the embedding of gene expression for model input. Our approach for binning the expression may cause some resolution loss. Second, gene interactions usually exist in the form of networks (i.e., gene regulatory networks and biological signaling pathways)^43^, and this kind of prior knowledge hasn’t been explicitly incorporated in scBERT. Aggregating information from neighbors within a graph neural network based on biological networks may better mimic the gene-gene interaction. The idea could be applied to the single-cell analysis by building cell-level graph using the scRNA-seq data. From this point of view, it can be foreseen that Transformers for graph^44^ may be the future development direction of scBERT^45^. Third, the efficiency of masking during pre-training is another point worth optimizing. Current masking strategy in scBERT is simplified with non-zero masking. With the zero-inflated input^46^, the model might be inclined to output all zeros for the reconstruction task during pre-training. Therefore, we masked the non-zero values and calculated the loss based on the non-zero values during pre-training. However, masking only the non-zero values may lower the utilization of the single-cell data for pre-training, due to their minority. Advanced masking strategy tailored for single-cell data could be introduced to improve the computational efficiency of the masking process.

For future work, we would like to explore the versatility and flexibility of scBERT in a variety of downstream tasks (i.e., gene-gene interaction, batch correction, clustering, differential analysis in disease conditions)^47^.

## Methods

### The scBERT model

The scBERT model adopts the advanced paradigm of BERT and tailor the architecture to solve single cell data analysis. The connections of our model with BERT are given as follows.

First, scBERT follows BERT’s revolutionary recipe to conduct self-supervised pretraining^25^ and to use Transformer as the model backbone^33^.

Second, our design of embeddings is similar to BERT in some aspects while having unique features to leverage gene knowledge. From this perspective, our expression embedding could be viewed as the token embedding of BERT. Since shuffling the columns of our input does not change its meaning (like the extension of BERT to understand tabular data with TaBERT^27^), absolute positions are meaningless for gene. Instead, we use gene2vec to produce gene embeddings, which could be viewed as relative embeddings^26^ that capture the semantic similarity between any of two genes.

Third, Transformer with global receptive field could effectively learn global representation and long-range dependency without absolute position information, achieving excellent performance on non-sequential data (i.e., images, tables)^24,27^.

#### Gene embedding

In NLP, the inputs of the BERT model are word embeddings, a set of real-valued vectors in a predefined vector space that represent individual words. The word embedding technology helps to better represent the text by assuring the words with similar meanings have a similar representation^47^. While from the aspect of scRNA-seq, the inputs are constituted by individual genes and a predefined vector space is in need for representing the similarity between genes. Hence we employed gene2vec^28^ to specifically encode gene embeddings. In this way, the difficulty of model training is reduced, with the help of the inter-gene relationship provided by prior knowledge.

#### Expression embedding

In spite of the gene embedding, there is also a challenge on how to utilize the transcription level of each gene, which is actually a single continuous variable. It’s worth noting that the frequency of the word occurrence in a text is valuable information for text analysis and is often transformed as bag-of-words by term frequency statistic analysis for downstream tasks in the area of NLP^48^. The gene expression can also be considered as the occurrence of each gene that has already been well-documented in a biological system. From this insight, we applied the conventionally-used term frequency analysis method that discretizes the continuous expression variables by binning and converts them into 200-dimensional vectors, which are used as token embedding for the scBERT model.

#### Model Building

The quadratic computational complexity of the BERT model with Transformer as the basic unit does not scale very well to long sequence while the gene number of scRNA-seq can be up to more than 20,000. To this end, a matrix decomposition version of the Transformer, Performer was employed to enlarge the sequence length. The regular dot-product attention in Transformer is a mapping of *Q, K, V*, which are encoded representations of the input *queries, keys*, and *values* created for each unit respectively. The bidirectional attention matrix is formulated as:

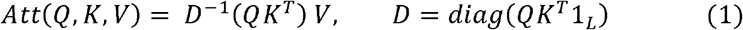

where *Q* = *W_q_X, K* = *W_K_X, V* = *W_V_X* are linear transformations of the input *X*, and *W_Q_, W_k_, W_v_* are the weight matrices as parameters. 1*_L_* is the all-ones vector of length *L*, and *diag*(·) is a diagonal matrix with the input vector as the diagonal.

The attention matrix in Performer is described as follows:

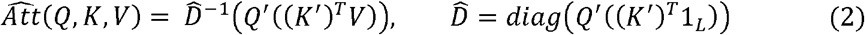

where *Q*′ = Ø(*Q*), *K*′ = Ø(*K*), and the function Ø(*x*) is defined as:

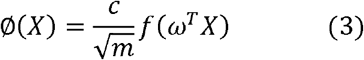

where *c* is a positive constant, *ω* is a random feature matrix, and *m* is the dimesionality of the matrix.

Here, we constructed our model with 6 Performer enocoder layers and 10 heads for each layer.

The model training process contains two stages: self-supervised learning on unlabeled data to get a pre-trained model and supervised learning on the specific cell type annotation tasks to get the fine-tuned model.

#### Self-supervised learning on unlabeled data

In this study, we followed the conventional self-learning strategy of the BERT model in NLP tasks by randomly masking the input data value and making a prediction based on the remaining inputs. Considering the dropout zeros phenominan^49^, we randomly masked the non-zero gene expression and then reconstructed the original inputs by model predictions using the remaining genes. We ultilized cross-entropy loss as the reconstruction loss, formulated as:

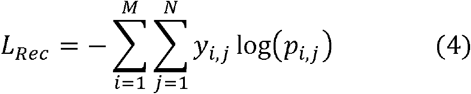

where *M* is the number of cells, *N* is the number of masked gene expression values, while *y_i,j_* and *p_i,j_* are the true expression and predicted expression of gene *j* in cell *i* respectively. Through this self-supervised strategy, the model can learn general deep representations of gene expression patterns on the large amount of unlabeled data which might alleviate the efforts of the downstream fine-tuning process.

#### Supervised learning on specific tasks

The output of scBERT was a 200-dimension feature corresponding to each gene and a one-dimensional convolution was applied for abstract information extraction for each gene feature. Then a 3-layer neural network was applied as the classification head and transformed the gene features into the probability for each cell type. Cross-entropy loss was also employed as the cell type label prediction loss, calculated as:

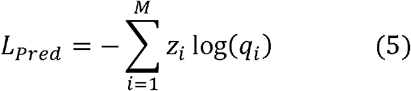

where *z_i_* and *q_i_* indicates the ground truth cell type label and predicted label of cell *i*.

### Datasets

Since the model training consists of two stages: self-supervised learning on unlabeled data and fine-tuning on task-specific data, the dataset used in the two stages were collected from different sources to avoid data leakage. In the first stage, large amounts of data without annotations were used for general pattern learning. In the second stage, task-specific data with well-annotated cell labels were required for the subsequential systematic benchmarking of scBERT and SOTA methods. To this end, we only included scRNA-seq datasets that provided highly credible cell type annotation and had been cited by the majority of the cell type annotation methods for performance evaluation.

#### The Panglao dataset

The Panglao dataset^50^ was downloaded from the PanglaoDB website (https://panglaodb.se/). In brief, PanglaoDB integrated 209 human single-cell datasets consisting of 74 tissues, and 1,126,580 cells that originated from different experiment sources via various platforms. In this study, we used scRNA-seq data from PanglaoDB for first stage pre-training. No annotations or cell labels were used at the first stage since the self-learning strategy was employed and only the genes and their expression level were needed as inputs for the scBERT model.

#### Zheng68k dataset

The Zheng68k is a classic PBMC dataset by 10X CHROMIUM that is widely used for cell type annotation performance acessment^35^. It contains about 68, 000 cells within eleven subtypes of cells: CD8+ Cytotoxic T cells (30.3%), CD8+/CD45RA+ Naive Cytotoxic cells (24.3%), CD56+ NK cells (12.8%), CD4+/CD25 T Reg cells (9.0%), CD19+ B cells (8.6%), CD4+/CD45RO+ Memory cells (4.5%), CD14+Monocyte cells (4.2%), Dendritic cells (3.1%), CD4+/CD45RA+/CD25-Naive T cells (2.7%), CD34+ cells (0.4%), CD4+ T Helper2 cells (0.1%). Zheng68k dataset contains rare cell types and the distribution of cell types in this dataset is imbalanced. Meanwhile, strong correlations between cell types make it difficult to differentiate them.

#### Pancreas datasets

The pancreas datasets consist of Baron, Muraro, Segerstolpe, and Xin. The cell type labels were aligned and four cell types were included. The Baron dataset was downloaded from the Gene Expression Omnibus (GEO) (accession number: GSE84133), the protocol was inDrop^36^. The Muraro dataset was downloaded from GEO (accession number: GSE85241), the protocol was CEL-Seq2^37^. The Segerstolpe dataset was accessed from ArrayExpress (accession number: E-MTAB-5061), the protocol was Smart-Seq2^38^. The Xin dataset was downloaded from GEO (accession number: GSE81608), the protocol was SMARTer^39^. The above pancreas datasets were generated from different experiment platforms (Supplemental Table 1).

#### MacParland dataset

MacParland dataset^51^ from human liver tissue contains 20 hepatic cell populations from the transcriptional profiling of 8444 cells by 10X CHROMIUM. We downloaded the data from GEO (accession number: GSE115469) and generated the cell type annotation following the authors’ reported procedure.

#### Heart datasets

The heart datasets contain one large dataset^52^ for pre-training and Tucker dataset^53^ for benchmarking and evaluation in the hyperparameter sensitivity analysis. The large heart dataset for pre-training contains 451,513 cells from 11 cell types by 4 different sequencing platforms (Harvard-Nuclei, Sanger-Nuclei, Sanger-Cells, and Sanger-CD45) and was downloaded from the website (https://data.humancellatlas.org/explore/projects/ad98d3cd-26fb-4ee3-99c9-8a2ab085e737). The Tucker dataset contains 287,269 cells from 11 cell types via single nuclear RNA sequencing and was downloaded from the website (https://singlecell.broadinstitute.org/single_cell/study/SCP498/transcriptional-and-cellular-diversity-of-the-human-heart).

#### Lung dataset

The lung dataset was from human lung tissue and analyzed for COVID-19 related disease mechanism^54^. The dataset contains samples from 12 donors by 10X Genomics sequencing, and 39,778 cells from 9 cell types. The data was downloaded from the website (https://doi.org/10.6084/m9.figshare.11981034.v1).

#### Human Cell Atlas dataset

The Human Cell Atlas dataset^55^ contains 84,363 cells from 27 cell types among 15 major organs (skin, esophagus, trachea, heart, spleen, common bile duct, stomach, liver, blood, lymph node, small intestine, bladder, rectum, marrow, muscle) by HiSeq X Ten sequencing. The dataset was downloaded from GEO (accession number: GSE159929).

### Data preprocessing

As for the data provided in gene expression matrix format, log-normalization using a size factor of 10, 000 and quality control by filtering cell outliers with less than 200 genes expressed were performed on the data. As for the input of scBERT, no dimension reduction or HVG selection was processed since scBERT has a capacity of more than 20,000 genes as input and retains full gene-level interpretability.

### Comparison methods

For benchmarking, we implemented SOTA methods from the three categories, marker-based, correlation-based and supervised classification. Among them, SCINA, Garnett, and scSorter stand for the annotation using marker gene databases; Seurat, SingleR, CellID, and scmap are correlation-based methods; scNym and Scibet are the SOTA methods that conduct annotation by supervised/semi-supervised classification. Notably, this categorization depends on how the most important process is conducted. As for marker gene-based annotation, CellMarker database with manually curated cell-type markers using a literature search of over 100 000 papers was applied for the marker database^56^. No manual selection of the marker genes was included for an unbiased and fair comparison of all the methods.

#### scNym

scNym is a recently proposed semi-supervised learning annotation method that leverages the unlabeled target data through training a domain adversary^57^. It requires no prior manual specification of marker genes. It makes use of the target data by domain adaptation and achieved the SOTA performance on several tasks. However, users have to endure the inconvenience that they must re-train the model on each batch of the new coming data.

#### SciBet

Scibet is a supervised classification method that selects genes using E-test for multinomial model building and annotates cell types for a new cell in the test set^29^. We adopted SciBet in R package for benchmarking.

#### Seurat

As a popular single-cell data analysis pipeline, Seurat is widely used by biologists and clinical experts. Seurat maps the query samples to the reference dataset in a reference-based annotation manner^58^. In this study, we adopted the implementation of the cell type annotation of Seurat V4.0 and followed the cell type annotation tutorial provided by Seurat for benchmarking.

#### SingleR

SingleR is a reference-based analysis method that calculates the Spearman coefficient on variable genes and aggregates the coefficients to score the cell for each cell type^59^. It iterates the above process by subsampling top genes until the most closely related cell types are distinguished. The SingleR package was applied for benchmarking.

#### CellID

CellID is a clustering-free multivariant statistical method for cell type annotation that performs dimensionality reduction, evaluates gene-to-cell distance, and extracts gene signatures for cells (cell-to-cell strategy) and groups (group-to-cell strategy)^30^. In this study, both strategies from the R package were used for benchmarking.

#### scmap

scmap is a reference-based annotation method including two strategies: scmap_cluster and scmap_cell. scmap_cluster maps individual cells from query samples to certain cell types in the reference dataset while scmap_cell maps individual cells from query samples to individual cells in a reference dataset^31^. Both scmap_cluster and scmap_cell perform feature selection and calculate the distance (cosin and euclidean distance). The reference is searched for the nearest neighbors to a query cell. We used the R package of scmap for the scmap-cluster and scmap_cell tools.

#### SCINA

SCINA is a typical marker gene-based annotation method that requires a list of marker genes for different cell types and identifies the cell types based on the assumption that there exists a bimodal distribution for each marker gene and the higher modes belong to the relevant cell type^9^. We used the Scina package for benchmarking.

#### Garnett

Garnett requires a user-defined cell hierarchy of cell types and marker genes as input. Garnett aggregates marker gene scores using term frequency-inverse document frequency transformation and uses an elastic-net regression-based model for annotation^10^. We adopted the original R package to use the garnet model for benchmarking.

#### scSorter

Scsorter employs marker genes and the HVGs for clustering and cell type annotation based on the observation that most marker genes do not consistently preserve high expression levels in all the cells that belong to the related cell types^32^. Here we adopted the R implement of Scsorter.

### Benchmarking

To assess the performance of the annotation methods under different scenarios, 9 pairs of reference and test datasets were generated and the performance was evaluated using scBERT and all the above methods. The details were listed below.

#### Performance on intra-dataset data using cross-validation

The PBMC data from Zheng68k with high inter-class similarity, the Pancreas datasets (Baron, Muraro, Segerstolpe, and Xin), the MacParland dataset, the Tucker dataset, the Lung dataset,t and the Human Cell Atlas dataset were employed for testing the performance on the intra-dataset in a 5-fold cross-validation manner. Notably, the reference dataset in this section also refers to the training dataset for the supervised methods, including scBERT.

#### Performance on the inter-dataset data

To evaluate the robustness of the methods on cross-cohort data with batch effects from different single-cell sequencing platforms, we tested the methods on four pancreas datasets (Baron, Muraro, Segerstolpe, and Xin), taking three datasets as the training set and the remaining one as the test set each time. Considering the difference in cell populations among these datasets, all datasets were aligned, retaining only four kinds of pancreas islet cells (alpha cell, beta cell, delta cell, and gamma cell) that are common in these datasets. To evaluate the robustness of the methods on cross-organ data, we tested the methods on three major organ (esophagus, rectum an stomach) from Human Cell Atlas dataset.

#### The influence of reference cell amount on the performance

The number of reference cells is prone to influence the model performance. In this study, 10%, 30%, 50%, 70%, and 90% of the PBMC cells from the Zheng 68K dataset were randomly selected as the reference for finetuning while the remaining as the query samples for testing.

#### Class-imbalanced data tests

Following the construction method for class-imbalanced data^4^, we collected four PBMC cell types, CD19+B, CD8+Cytotoxic T, CD34+, and CD8+/CD45RA Naïve Cytoxic cells that contain various levels of similarity across cell types from Zheng68K data. The cells of the four types were randomly selected with the cell numbers 10000, 100, 10000, and 100 as reference data for finetuning. As for model testing, 100 cells were randomly selected per cell type as query data.

#### Novel cell type detection

Human liver tissue was used to assess the unknown cell type identification. Here we adopted MacParland dataset^51^ from human liver tissues with 8,434 cells belonging to 14 cell types. In this experiment, we took four immune cells for novel cell type simulation, which were absent from other liver datasets. Following the schema proposed in the previous study^7^, we performed leave-out one cell type evaluation by removing one cell type from the reference dataset while keeping the cell type groups in the query dataset. The evaluation process was iterated on each cell type. At present, there is no unified quantitative evaluation metrics for detection of novel cell type. Some approaches compute the accuracy by putting the novel class together with known classes, which unavoidably overwhelms the models’ accuracy for rare and novel cell types. Besides accurately detecting novel cell types, a good cell type annotation method should maintain the ability to accurately discriminate known cell types. In this regard, we evaluate the accuracy of novel cell type and known cell types, separately. Notably, we employed a strict evaluation method for novel cell types with the accuracy calculated on the union set of cells with the novel cell type label and the cells that are predicted as novel cell types.

#### Assessment on the necessity of self-learning

To illustrate the necessity of the self-learning process of scBERT, the performance gain was evaluated on the model after self-learning and finetuning compared to the model training from scratch.

#### Evaluation metrics

Cell type annotation performance of each method at cell-level and cell-type-level was evaluated using the metrics of accuracy and macro F1 score, respectively. Since cell type annotation task and cell clustering task are not equivalent, those metrics assessing the quality and distance of clusters are excluded from this study.

#### Sensitivity analysis on the hyperparameters

The influence of the hyperparameters (size of the embedding vector, the binning setting, the number of encoder layers, and the number of heads for each layer) were systematically estimated on the heart datasets with large-scale heart dataset (451,513 cells) as the pre-training dataset and the Tucker dataset as the evaluation dataset.

#### Scalability

When evaluating on the large Tucker datasets with 287,269 cells, those comparison methods implemented in R faced severe problem in scalability due to their poor memory management. For instance, CellID met the memory bottleneck when calculating a matrix of 50,000 × 230,000 and we made efforts to split the matrix into pieces to avoid memory overflow. Conversely, benefiting from mini-batch sampling and the efficient Performer encoder, scBERT could easily deal with large-scale datasets at both the pre-training and the fine-tuning stage.

#### Marker genes for the marker-based comparison methods

To avoid bias introduced by marker selection, well-documented marker lists associated with well-defined cell types from CellMarker^56^ were used.

### Systematic analysis of scBERT

#### Pre-training versus not pre-training

Following the BERT’s pre-training and fine-tuning paradigm, our method is prone to generate an efficient encoder and provide a general embedding that better represents the gene expression of each cell by revealing critical patterns with lower data noise. The results of ablation study on model performance with and without pre-training (Extended Data Figure 1a) proved the essentiality of pre-training for the model’s down-streaming task (i.e., cell type annotation) with a relatively large and significant difference in the bioinformatics field. The scBERT model extracts the useful attention pattern on gene expressions and interactions from a large scale of various scRNA-seq data, alleviating the efforts of the fine-tuning process on the specific downstream tasks.

#### Feasibility on classifying with gene expression patterns

It’s well known that marker genes play a key role in cell type annotation for marker gene-based annotation and most of the reference-based annotation. Even some of the supervised-based methods are heavily dependent on the prior marker gene knowledge. Among the current mainstream methods that use marker genes for classification, some methods utilize the gene expression pattern for cell type annotation. Both kinds of methods were reported to achieve good performance on variable cell type annotation tasks, indicating that both types of data imply discriminative information for different cell types. To investigate the effect of marker genes and the discriminant ability of remaining expression patterns consisting of only the non-marker genes, we conducted experiments where marker genes were eliminated gradually, leaving the remaining expression profiles for cell type annotation (Extended Data Figure 1b, Supplementary Table 16). The results proved that the marker genes are important for the cell type annotation. However, in addition to the marker genes, there are still informative gene patterns that have good distinguishing power on cell type classification. With deletion of 100% marker genes, scBERT can still efficiently learn the informative gene patterns and achieve the performance on par with the best performance achieved by comparison methods with all marker genes on the representative Zheng68K dataset (Extended Data Figure 1b). We also explored detected gene lists from scBERT and other ML method (i.e., scNym) and non-ML method (i.e. Seurat) on MacParland and Baron, respectively (Supplemental Table 17-18). Consistent with above experiment on the deletion of markers, we could observe that ML-based methods tend to learn high-level implicit cell-type-specific patterns (i.e. discovering some genes with high rank across cell types), whereas non-ML based methods usually simply find differentially expressed genes using statistics analysis. The results indicated that attention mechanism, saliency mechanism, and statistics analysis could gain complementary information from different perspectives on mining pattern of single-cell data.

#### General gene embedding vs. single-cell specific embedding

Gene2vec is based on bulk data^28^, which measures the average expression of genes from tissues and is the sum of cell type-specific gene expression weighted by cell type proportions^60^. In this regard, gene2vec maintains the general co-expression patterns of genes but stays away from strong noise and high sparsity of single-cell sequencing. Therefore, we utilized gene2vec as our gene embedding to represent the gene identity (each gene has a unique gene2vec embedding) and the semantic similarity from the aspect of general co-expression pattern. The encoder of scBERT could also learn single-cell specific embedding (we briefly call it scBERT embedding) that represents the cell-specific expression. To illustrate the evolvement of the embedding (or representation) during the model learning, we visualized the examples of gene2vec embedding and the scBERT embedding in Extended Data Figure 1b. Our model could generate different representations of the same gene for different cell inputs, while gene2vec generated all the same representation of the same gene for different cell inputs. We observed that the scBERT embedding exhibits cell-type-specific representation (the example representation of gene is significantly enriched in alpha cell type), which is naturally suitable for down-streaming cell type annotation task. Furthermore, the cell-type specific representation learns some correlation beyond gene2vec. Benefiting from the attention mechanism of the Performer, the model could detect the subtle gene interaction patterns that can only be seen in single-cell data after model training on scRNA-seq data (Extended Data Figure 1d). It could be observed that some genes have strong attention weights to all other genes, indicating that it plays a critical role in identifying the implicit patterns, which is consistent with the conclusion of the detected gene lists in Supplementary Table 17-18.

#### Influence of hyperparameters

A systematic investigation of the sensitivity of hyperparameters including the number of bins, the size of scBERT embedding vector, the number of attention heads, and the number of Performer encoder layers was performed on scBERT (Extended Data Figure 1b). First, the expression embedding by ranking the raw expression into 7 bins is suitable for scBERT. Increasing the bin numbers to 9 hampers the model performance, indicating that ranking the gene expression would denoise the raw data and improve scBERT’s efficiency in learning meaningful patterns. In the contrast, reducing the bin numbers would also affect the model performance, due to the loss of gene expression information (i.e., blurring the relatively large gene expression difference). The above experimental results proved that the proper design of bin number that balances the denoising and reserving expression information would benefit the model performance. Second, the gene2vec provided an embedding of 200 dimensions and achieved the best performance compared to other dimensions. Reduction of the dimension of scBERT embedding vector in the latent space would impair the model’s representation ability and performance (especially when the dimension is 50). Third, the Performer with 10 attention heads is suitable for our method. Decreasing the number of attention heads might reduce the model representation ability due to fewer representative subspaces. Increasing the number of attention heads seems to have limited influence on the performance. However, the over-parameterized model (with 20 attention heads) faces a risk of over-fitting, especially when applying to small datasets. Similarly, the model performs stable with 4 and 6 of Performer encoder layers but might suffer from under-fitting or over-fitting problem when decreasing or increasing the number of layers. Overall, the small fluctuations of the above parameters had little effect on the performance of the model, which also verified the robustness of scBERT.

### Model interpretability

We conducted a comprehensive interpretability analysis to explore the key genes for decision-making since scBERT models were built on the self-attention mechanism and all genes’ representation remained at the end of our workflow. The attention weights reflect the contribution of each gene and the interaction of gene pairs. The attention weights can be obtained from equation (1) modified by replacing *V* with *V*^0^, where *V*^0^ contains one-hot indicators for each position index. We integrated all the attention matrices into one matrix by taking an element-wise average across all attention matrices in multi-head multi-layer Performers. In this average attention matrix, each value *A*(*i,j*) represented how much attention from gene *i* was paid to gene *j*. In order to focus on the importance of genes to each cell, we summed the attention matrix along with columns into an attention-sum vector, and its length is equal to the number of genes. In this way, we could obtain the top attention genes corresponding to a specific cell type compared to other cell types. The attention weights were visualized and the top genes were sent to Enrichr^33^ for enrichment analysis.

Enrichment analysis was performed for the top-50-attention gene lists using various gene-set libraries and the results revealed there were some interesting relationships between top-enriched terms and the corresponding cell types.

### Statistical analysis

The Wilcoxon test was applied for the significance test. Cross-validation was employed in all the benchmarking experiments and standard deviations were drawn in the figures. The significance was calculated by wilconxon test on the paired groups. Jaccard index was used for similarity measure for the detected gene lists by different methods. Adjusted Rand Index (ARI) was applied to for similarity measure for clusters.

## Data Availability

All data used in this study are publicly available and the usages are fully illustrated in the Method section. The published Panglao dataset was downloaded from the PanglaoDB website (https://panglaodb.se/). The published Zheng68k dataset was downloaded from the “Fresh 68K PBMCs” on the website (https://support.10xgenomics.com/single-cell-gene-expression/datasets (SRP073767))^35^. The published pancreatic datasets were downloaded from the github (https://hemberg-lab.github.io/scRNA.seq.datasets/ (Baron: GSE84133, Muraro: GSE85241, Segerstolpe: E-MTAB-5061, Xin: GSE81608))^36–39^. The MacParland dataset was downloaded from the repository (https://www.ncbi.nlm.nih.gov/geo/ (GSE115469))^51^. The heart datasets were downloaded from the websites (https://data.humancellatlas.org/explore/projects/ad98d3cd-26fb-4ee3-99c9-8a2ab085e737 and https://singlecell.broadinstitute.org/single_cell/study/SCP498/transcriptional-and-cellular-diversity-of-the-human-heart)^52,53^. The lung dataset for COVID-19 study was downloaded from the website (https://doi.org/10.6084/m9.figshare.11981034.v1)^54^. The adult human cell atlas of 15 major organs dataset was downloaded from the website (https://www.ncbi.nlm.nih.gov/geo/ (GSE159929))^55^.

## Code Availability

The source code of the preprocessing, scBERT modeling, and fintuning processes are freely available on Github (link: https://github.com/TencentAILabHealthcare/scBERT, DOI: 10.5281/zenodo.6572672)^61^ with detailed instructions. The source codes of the other comparison methods are publicly available (web links and package versions are listed in Supplementary Table 2).

## Acknowledgements

We thank Biaobin Jiang and Yan Ji for their valuable suggestions on model building and experimental design. We thank Tao Shen for advice on the large-scale model pre-training. H.L. was supported by the National Key R&D Program of China [2018YFC0910500]; SJTU-Yale Collaborative Research Seed Fund; the Neil Shen’s SJTU Medical Research; Key-Area Research and F.Y. was supported by Development Program of Guangdong Province (2021B0101420005).

## Author Contributions

F.Y. and J.Y. conceived and designed the project. W.W. developed and implemented the algorithms under the guidance of F.Y. and J.Y. W.W. and F.W. collected the datasets. W.W., F.Y., and F.W. conducted the experiments, data analysis and method comparison. F.Y. and W.W.completed the figures and manuscript with the guidance of J.Y. and H.L. Y.F. and F.W. polished the manuscript and figures.

D.T. gave suggestions on the design of transformer architecture and applying NLP technology. J.H. gave suggestions on improving the manuscript. F.Y. and F.W. revised the figures and manuscript. All authors reviewed and approved the manuscript. F.Y., W.W., and F.W. contributed equally.

## Competing Interests

The authors declare no competing interests.

## Notes

### Competing Interest Statement

The authors have declared no competing interest.

### Summary of Updates

Update the abstract, author information, and the order of sections.

