## Supplementary figures and images for "scBERT as a Large-scale Pretrained Deep Language Model for Cell Type Annotation of Single-cell RNA-seq Data"

### Extended Data Figure 1

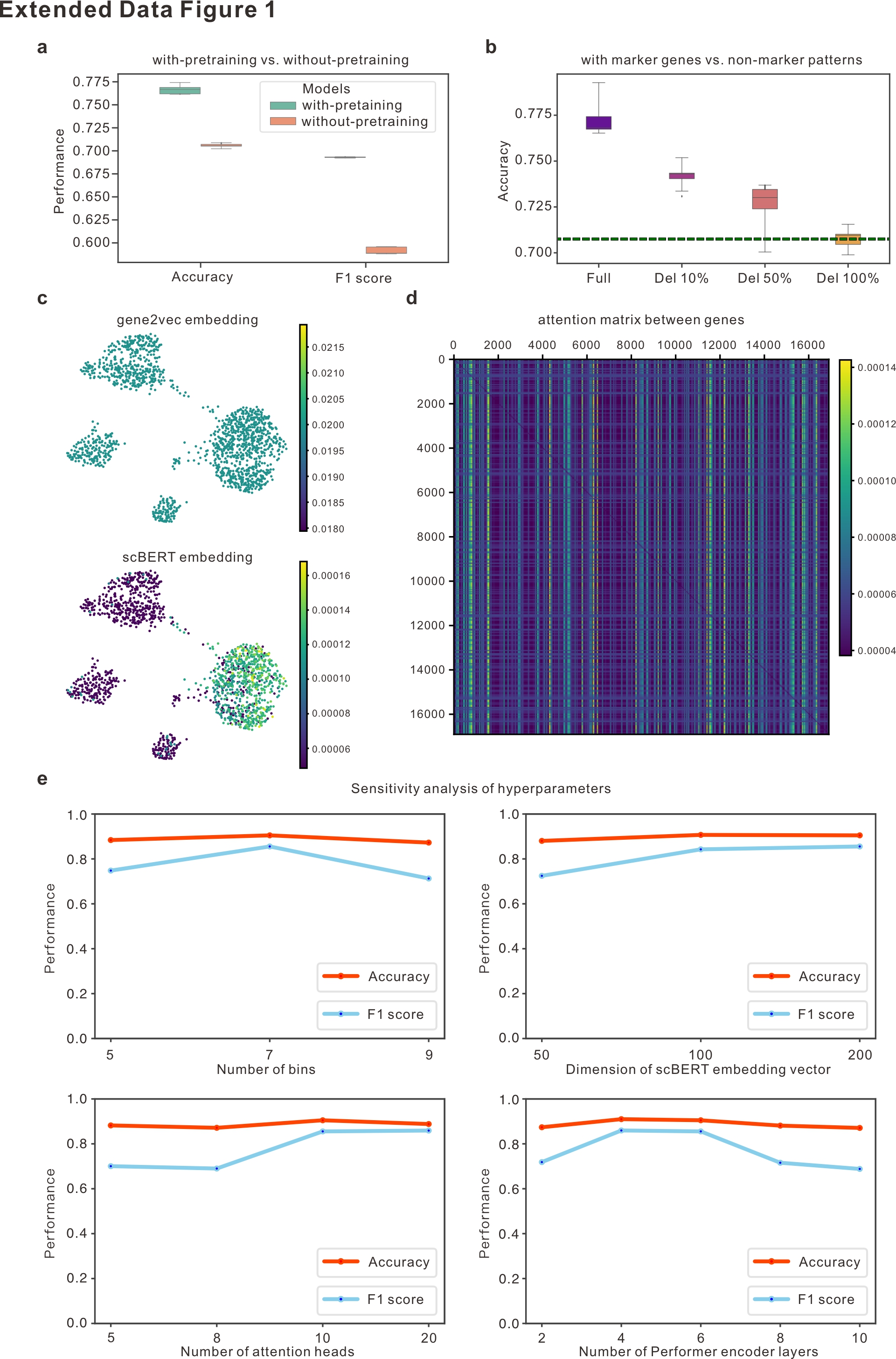

### Extended Data Figure 2

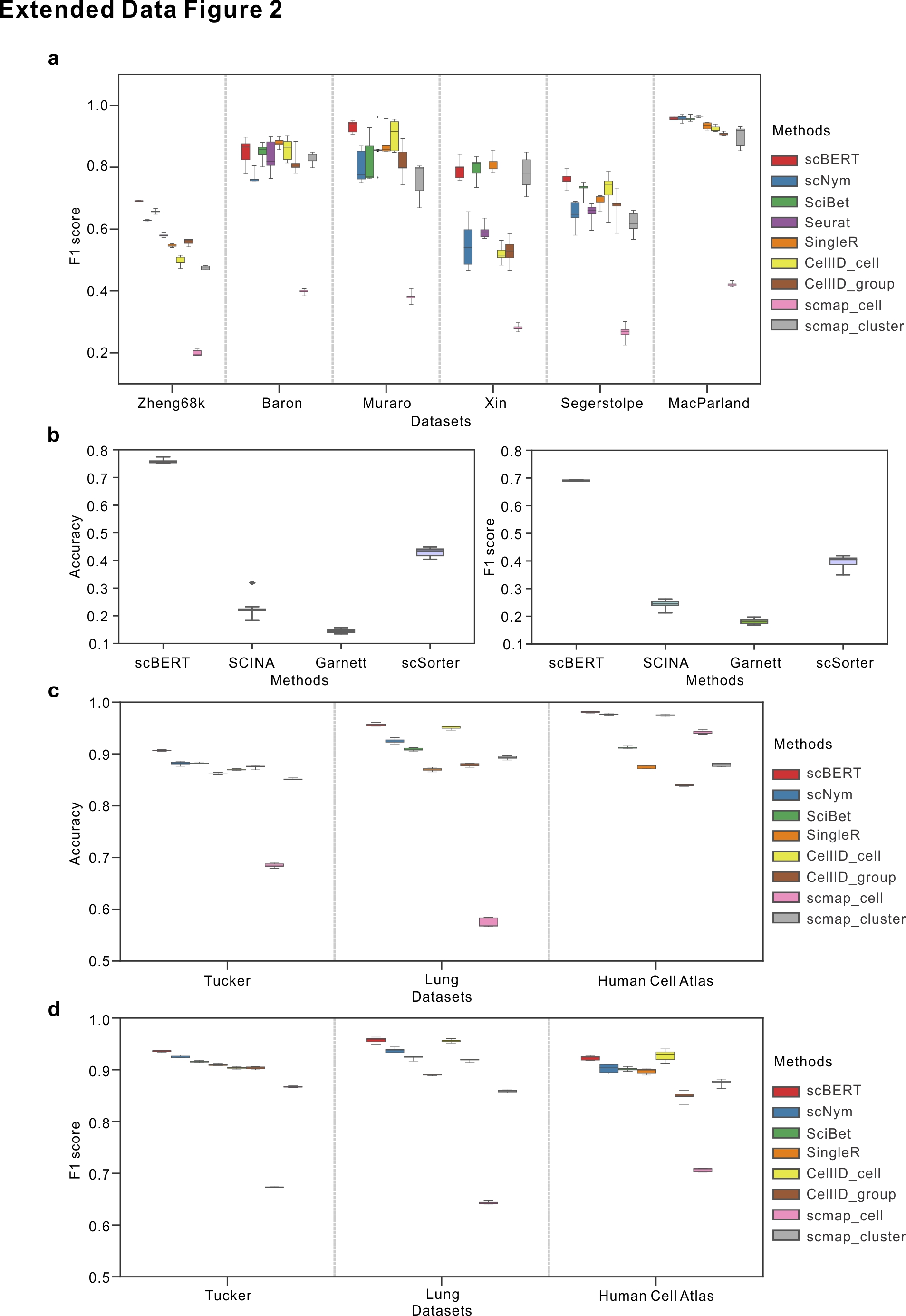

### Extended Data Figure 3

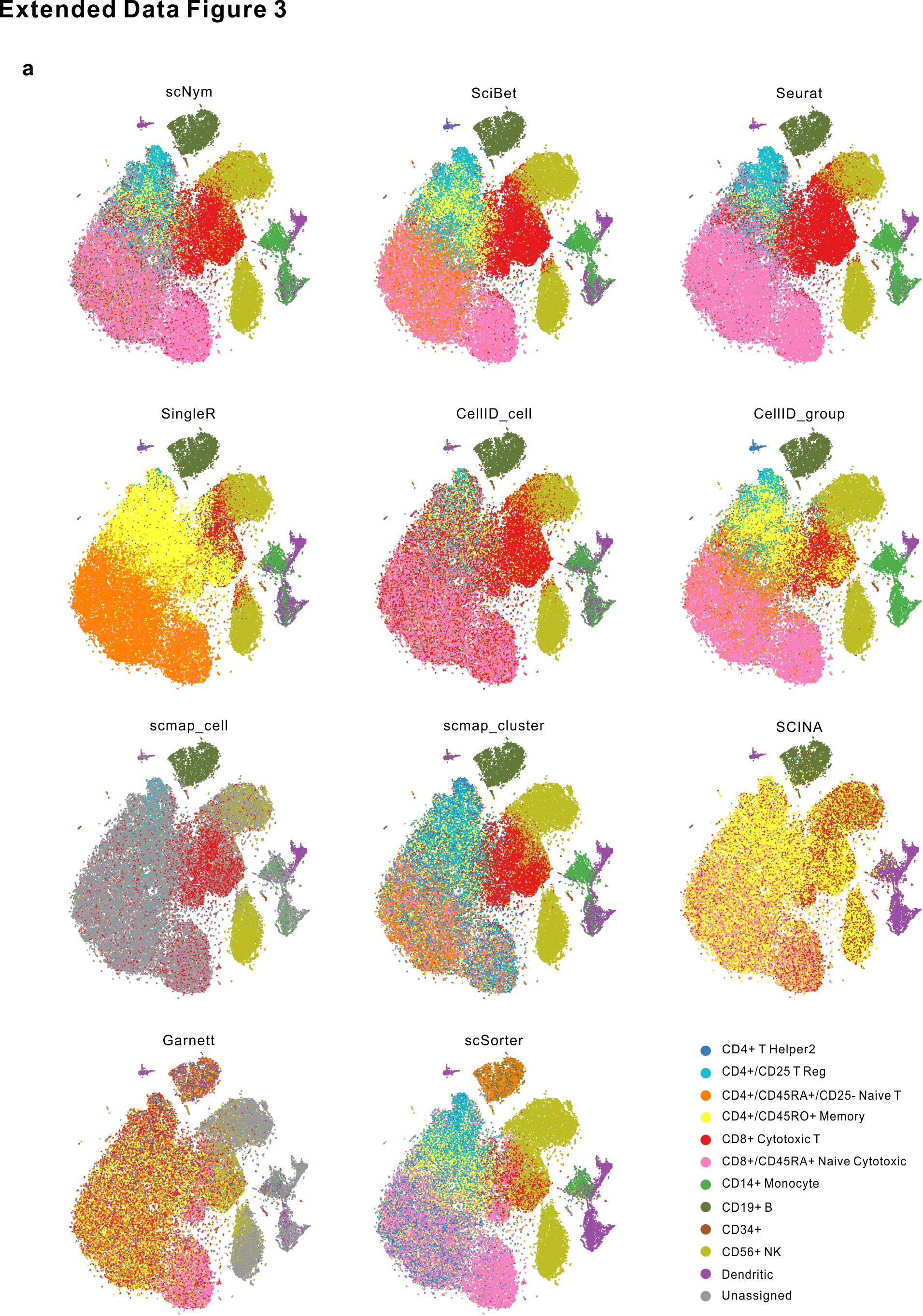

### Extended Data Figure 4

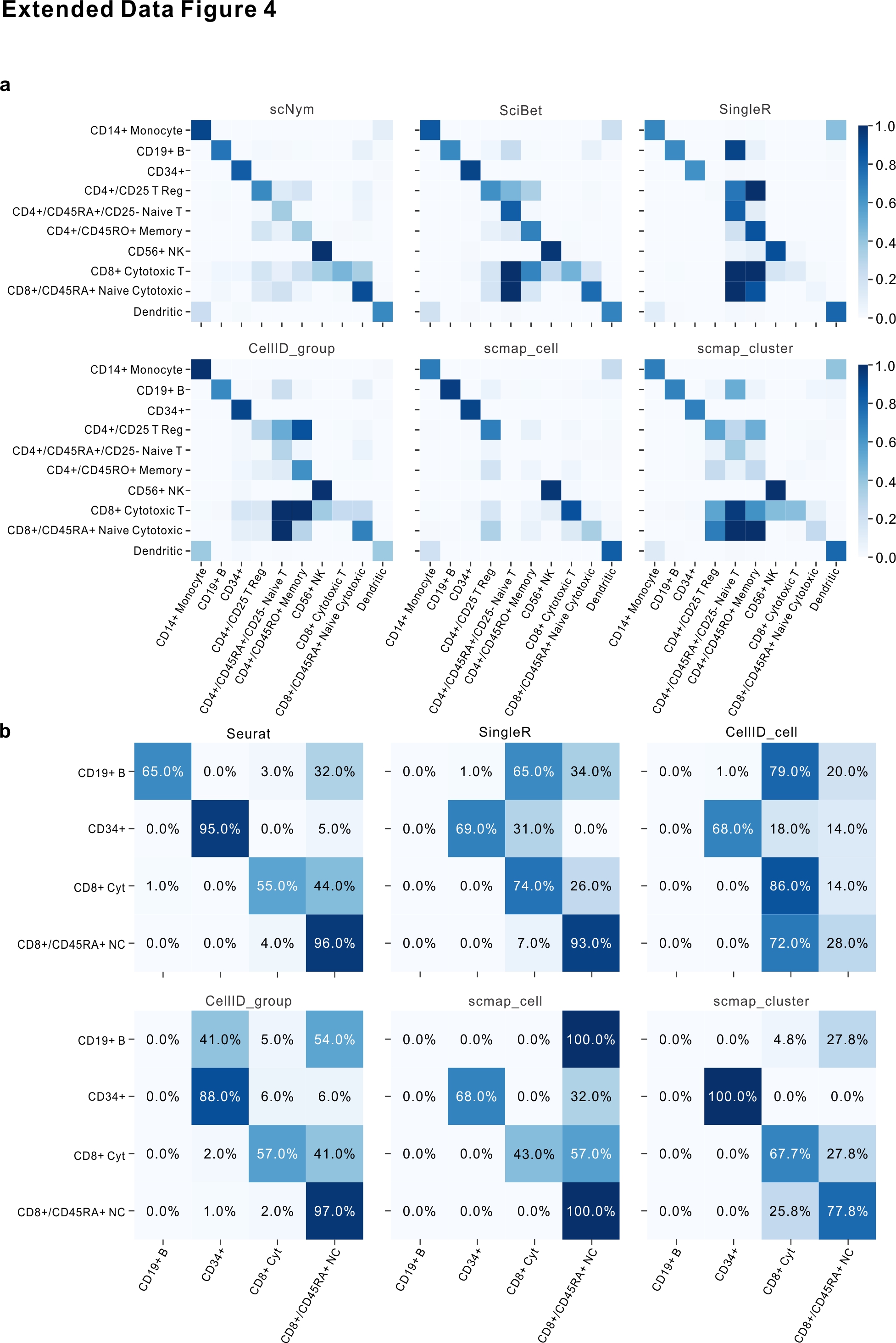

### Extended Data Figure 5

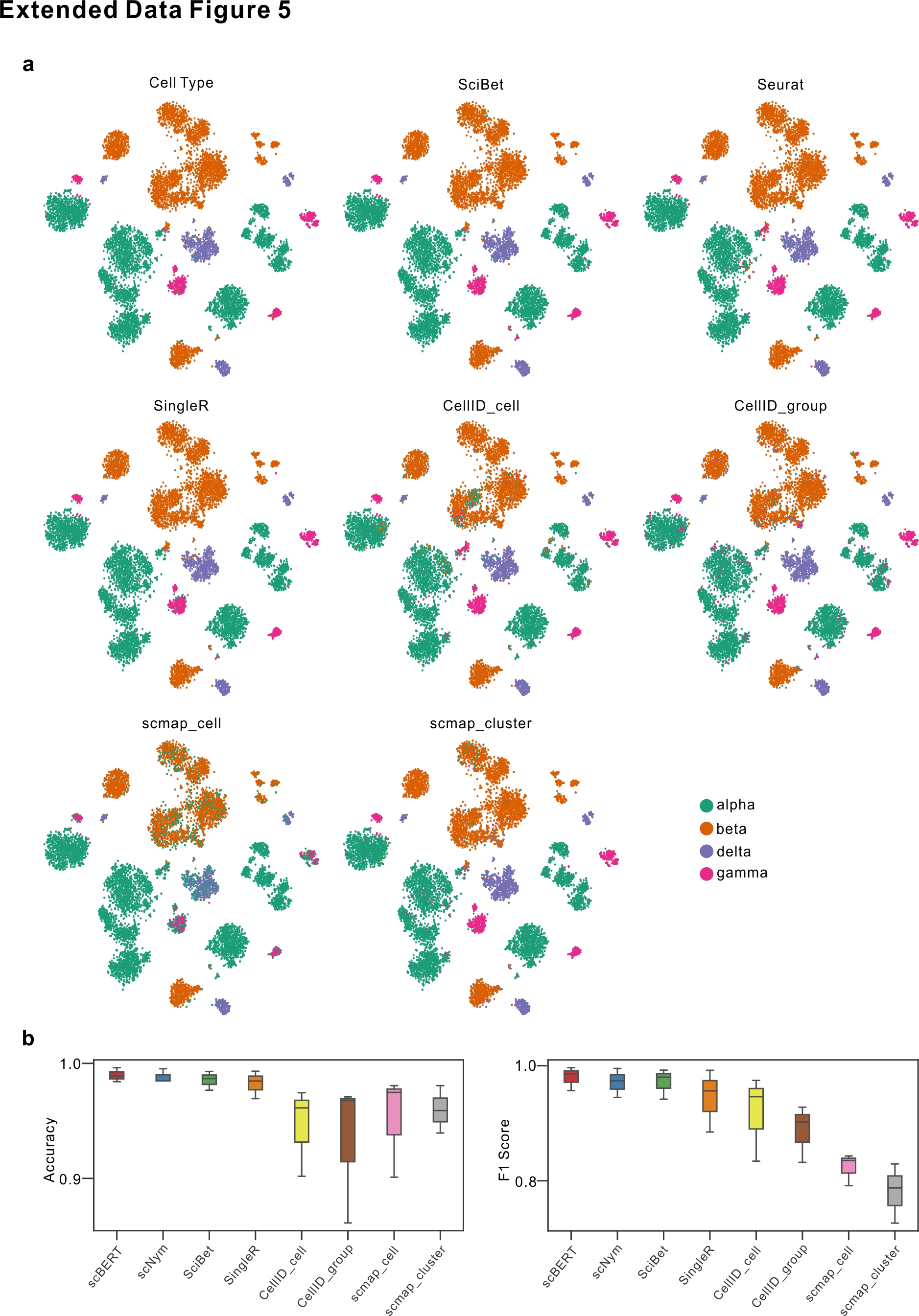

### Extended Data Figure 6

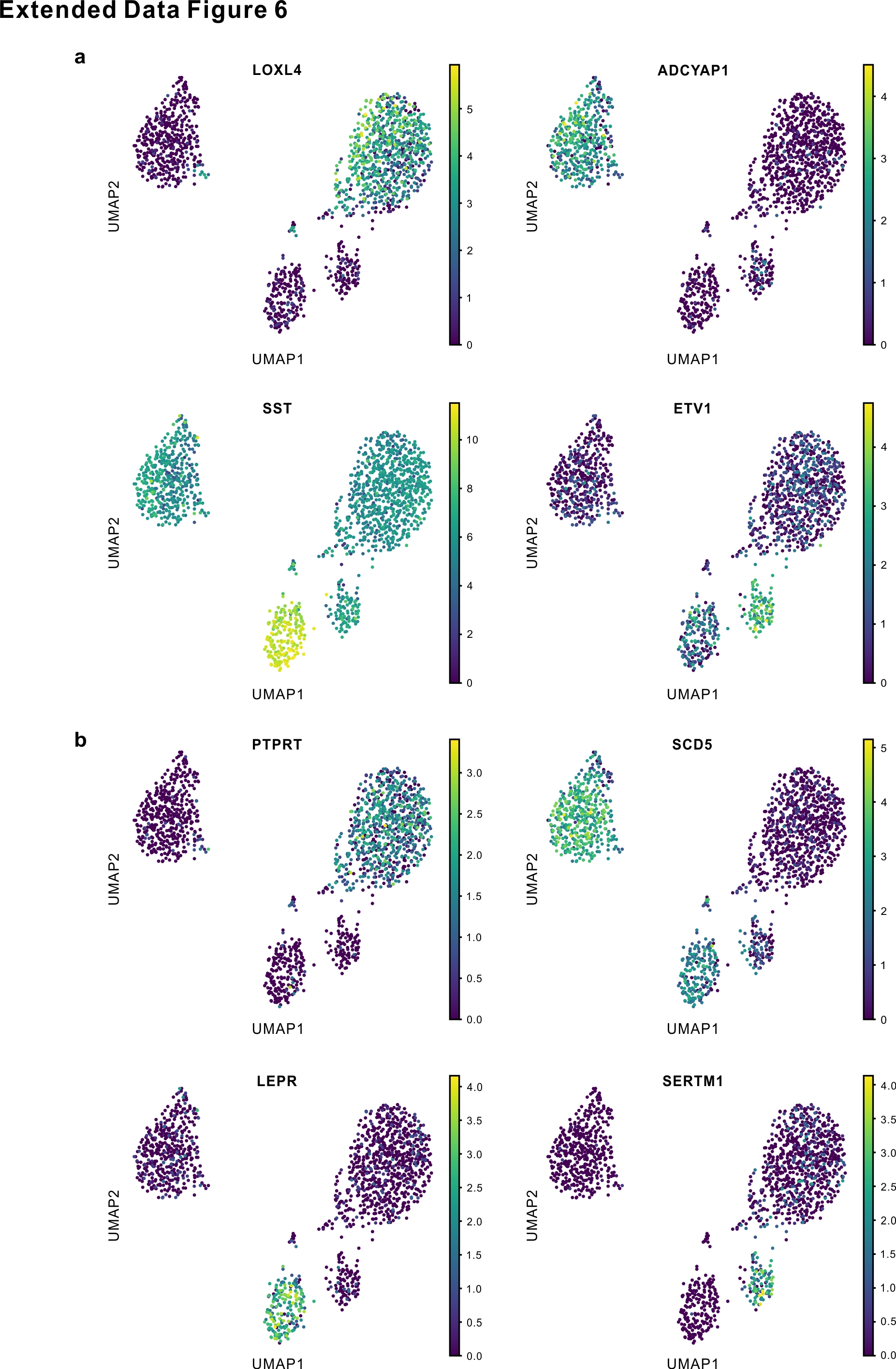
